# Spatial analysis of HPV associated cervical intraepithelial neoplastic tissues demonstrate distinct immune signatures associated with cervical cancer progression

**DOI:** 10.1101/2024.10.02.611623

**Authors:** Gianna Pavilion, Hani Vu, Zherui Xiong, Thi Viet Trinh Dang, Michael Walsh, Andrew Causer, Janin Chandra, Quan Nguyen, Ian H Frazer

## Abstract

Cervical cancer remains the fourth most common cancer affecting women worldwide, and incidences of other HPV-related cancers continue to rise. For the development of effective prevention strategies in high-risk patients, we aimed to better understand the roles of inflammatory pathways and the tumour microenvironment as the main driver of progression to malignancy in HPV-infected tissues. We analysed the spatial organisation of seven samples of HPV+ high-grade squamous intraepithelial lesion (HSIL) and cervical intraepithelial neoplasia 3 (CIN3), comparing tumour heterogeneity and immune microenvironments between pre-malignant (neoplastic) and adjacent cervical tissues. We observed evidence of immune suppression within the neoplastic regions across all samples and identified distinct immune clusters for each dysplastic lesion. Previous single-cell data analyses in an HPV16 E7 oncoprotein-driven transgenic mouse model suggested a potential role for IL34-CSF1R signalling in immune modulation, where low IL34 expression was associated with Langerhans cell dysfunction, and, in cervical cancer, with poor patient outcome. Here, we observed that IL34-CSF1R co-expression was absent within HPV-associated neoplastic tissues but present in adjacent normal tissue regions. Additionally, we identified enrichment of an M2 gene signature in neoplastic tissue, while adjacent tissue was enriched with a pro-inflammatory M1 gene signature. Our findings provide bio-pathological insights into the spatial cellular and molecular mechanisms underlying HPV-associated cervical cancer immune regulation and suggest a strategy to modulate the immune system in HPV-positive neoplastic cervical and other tissues.

## Introduction

Despite significant advances in biomedical research and public health interventions, cervical cancer persists as a major global health concern, ranking as the fourth most prevalent cancer among women worldwide (Brisson and Drolet 2019; Singh et al., 2020). Moreover, the incidence of human papillomavirus (HPV)-driven anogenital and oropharyngeal cancers is on the rise globally, affecting both men and women (Malagon et al., 2024; Lundberg et al., 2011; Rettig et al., 2018). Traditional histopathological examinations of tissue sections, while informative, lack the resolution to provide a comprehensive understanding of the complex genetic and immune-spatial landscape of cervical lesions. Current knowledge of CIN3 morphology primarily relies on insights derived from the study of E6 and E7 oncoproteins expressed by oncogenic high-risk HPV types (Cattani et al., 2009; Zhang et al., 2018; Singini et al., 2023; Peng et al., 2024). There remains much ambiguity surrounding the mechanisms underlying HPV persistence, immune dysregulation, and the progression from low-to high-risk intraepithelial lesions (Wang et al., 2018). Thus, a deeper understanding of these mechanisms is crucial for the development of effective prevention and treatment strategies.

One potential mechanism of immune suppression involves the dysregulation of cytokine signalling pathways that control immune cell recruitment and function. In particular, the interactions between Interleukin-34 (IL34) and its receptor, Colony Stimulating Factor-1 Receptor (CSF1R), have emerged as key modulators of immune responses in various cancers (Cersosimo et al., 2024; Otsuka et al., 2021). CSF1R is expressed exclusively on mononuclear phagocyte cells (Hume et al., 2020). IL34 has been identified as an alternative ligand for CSF1R, promoting monocyte survival (Lin et al., 2008). While IL34 is expressed in various tissues, it is most abundantly found in the skin and brain, where it plays a crucial role in the maintenance and development of Langerhans cells (LCs) and microglia, respectively (Greter et al., 2012; Wang et al., 2012). IL34-CSF1R signalling can lead to activation of various pathways, including STAT, AKT, and ERK1/2, caspase and autophagy, which in turn trigger cell proliferation, differentiation, migration, and cytokine production (Baghdadi et. al., 2018).

The role of IL34 in cancer progression is not fully understood, and whether it has a protective or detrimental effect depends on the tissue and cancer types (Lelios et. al, 2020). However, in the progression of cervical intraepithelial lesions to invasive cancer, our research identified a continuous decline inIL34 which also correlated with poor patient prognosis (Tuong et al, 2021). We previously demonstrated in a transgenic mouse model of HPV16 E7-mediated epithelial hyperplasia that IL34-CSF1R interaction was absent between hyperplastic keratinocytes and LCs (Tuong et al., 2021). This absence was associated with an imbalance in LC cell states, characterised by a depletion of immune-stimulatory LCs and an enrichment of immune-inhibitory LCs (Tuong et al., 2021). In line with this, we have found that LCs of HPV16 E7 mediated epithelial hyperplasia are dysregulated in phenotype, maturation and function (Abd Warif., 2015; Chandra et al., 2016; Bashaw et al., 2019; Tuong et al., 2021).

These findings suggest that HPV-driven epithelial hyperplasia may impair LC function and contribute to immune evasion, potentially facilitating malignant progression (Tuong et al., 2021). Additionally, a mouse model described by Strachan et al. (2013), investigated the role of the CSF1R signalling pathway in recruiting tumour-associated macrophages (TAMs), revealing that CSF1R inhibition decreases TAM turnover and tumour growth while increasing T cell infiltration. This suggests continuous CSF1R blockade as a potential anticancer therapy. These murine models have been valuable in simulating the pathophysiology of high-grade squamous intraepithelial lesions (HSIL) and furthering our understanding of the role of LCs and TAMs in carcinogenesis (Strachan et al., 2013).

Examining the spatial distribution of immune cells and neoplastic epithelial cells within human CIN3 lesions can yield insights unattainable with traditional methods or murine models. The advent of spatial and multi-omics approaches offers unprecedented resolution and contextual depth for studying the tumour immune microenvironment (Causer et al., 2023). Recent single-cell analyses have uncovered key molecular signatures, specific signalling pathways, and distinct cellular communities within cervical cancer (Ou et al., 2022; Kurmyshkina et al., 2023; Guo et al., 2023; Liu et al., 2023; Qiu et al., 2023). However, a comprehensive spatial analysis of human HSIL/CIN3 samples is lacking.

In this study, we leverage spatial methodologies to conduct a detailed immuno-spatial analysis of human CIN3 samples. We explore ligand-receptor pair interactions and the co-expression of IL34 and CSF1R in HPV+ neoplastic human tissues and discuss the mechanistic parallels in relation to other HPV-driven cancers. Our findings reveal bio-pathological insights into the immune landscape of CIN3 and identify a potential alternative pathway for therapeutic intervention.

## Methods

### Ethics approval and sample selection

This study was approved by The University of Queensland Human Research Ethics Committee A (Project No. 2022/HE001601) and conducted in accordance with the National Statement on Ethical Conduct in Human Research (2007, current version). Samples were selected by pathologists at Sullivan Nicolaides Pathology (SNP) and corresponding p16 immunohistochemistry staining was performed on each biopsy to confirm the diagnosis of CIN3 (Sup Fig 1.). Ultimately, eight samples were selected on the criteria of no initial patient treatment and prominent p16 staining indicating abnormal cells affecting more than two thirds of the epithelium. Additionally, all lesions originated within the transformation zone of the cervix. Histopathological assessment was further conducted by a qualified pathologist on a matched reference H&E tissue section to determine a specific region of interest to analyse within a 6.5mm x 6.5mm Visium capture area.

### Tissue preparation - H&E and imaging

Eight 5-μm formalin-fixed, paraffin-embedded (FFPE) CIN3 samples, cut on SuperFrost slides, were provided by SNP. In accordance with the 10X Genomics Demonstrated Protocol (#CG000520), tissue sections were de-paraffinised and stained in Mayer’s haematoxylin (Dako) for 5 minutes and eosin (Sigma) for 1 minute. Imaging was performed using the Leica Aperio XT Brightfield Automated Slide Scanner with 20X magnification. Following imaging, sections were de-stained and de-crosslinked in preparation for library construction.

### Immunofluorescent staining

The immunofluorescent staining was performed sequentially for CD206, CD86 and p16 using the Opal-TSA system (AKOYA Biosciences). Briefly, adjacent CIN3 FFPE sections were deparaffinized with xylene and rehydrated through a gradient ethanol wash. Heat-induced epitope retrieval was performed using Target Retrieval Solution, Citrate pH 6 (Dako, S236984-2) at 100°C for CD206 and CD86; and with EDTA pH 9.0 at 105°C for p16. Background Sniper (Biocare Medical, BS966) was used as blocking buffer to minimize unspecific binding. Primary antibodies were sourced as follow: CD206 (Abcam ab64693); CD86 (Abcam ab243887), and p16 (BD Biosciences G1775-405). The TSA was applied using Opal 520 (AKO-FP1487001KT) for CD206, Opal 690 (AKO-FP1497001KT) for CD86, and Opal 570 (AKO-FP1488001KT) for p16.

### Library generation and sequencing

Sequencing libraries were constructed using the Visium CytAssist Spatial Gene Expression for FFPE, Human Transcriptome, 6.5mm kit (10X Genomics, PN #1000520) according to the Visium Spatial Gene Expression User Guide Rev C (10X Genomics, #CG000495). A total of 10-13 PCR cycles were employed for the amplification of the final Visium libraries. A qualitative assessment of each library was performed. Paired-end dual-indexing was performed on the NovaSeq 6000 platform (Illumina) using the following protocol; Read1 – 28bp, i7 – 10bp, i5 – 10bp, Read2 – 50bp. Illumina sequencing base call data (BCL) was demultiplexed using the Bcl2Fastq conversion software (v2.20). The SpaceRanger computational pipeline (10X Genomics, V2.0) was used to align the sequencing data to the GRCh38 human reference genome and map the data to the associated spatial coordinates of the H&E image, determined by spatial barcode information.

### Quality control, normalisation and batch correction

#### Quality control

Various processing issues during sample preparation such as tissue dissociation, folding, and other technical issues may lead to artefacts. These may translate into the data as variations in sequencing depth and batch effects. Mapped and aligned data was imported into the R package Seurat (v5.1.0) for data analysis. Genetic expression levels were determined based on the number of unique molecular identifiers (UMI). Outliers and low-quality reads were removed by filtering spots at a fixed threshold of < 100 counts per spot. Sparse genes expressed in < 3 spots were also removed. Sample VLP90_D was excluded from downstream analysis due to inadequate UMI counts.

#### Data normalisation

The Seurat *SCTransform* function was applied to normalise and scale filtered data. The function employs a regularised negative binomial regression, allowing biological variation to be retained within genetic expression profiles while eliminating technical variation and discrepancies among sequencing runs (Hafemeister & Satija, 2019). Preliminary selection of the top 3000 variable genes attained through this function was utilised for downstream analysis.

#### Batch correction and data integration

The normalised data were corrected for batch effect using Seurat’s *IntegrateData* function. The top 3000 variably expressed genes selected for each sample were leveraged to compute anchors through Canonical Correlation Analysis (CCA) and Mutual Nearest Neighbors (MNN) analysis, allowing technical differences to be mitigated (Stuart et al., 2019). To reduce dataset dimensionality, Principal Component Analysis (PCA) was conducted using the first 30 PCs. The number of PCs was selected based on elbow plot visualisation. Uniform Manifold Approximation and Projection (UMAP) embedding were generated to visualise the results.

### Clustering and Spot Type Annotation

Spatial transcriptomics (ST) techniques/technologies such as 10x Visium allow for deeper analysis of gene expression patterns whilst also preserving tissue morphology. Unsupervised clustering of the integrated dataset was performed with Seurat’s *FindClusters* function, where spots with similar gene expression profiles are grouped together using a shared nearest neighbour (SNN) modularity optimization based clustering algorithm (*FindNeighbors*). Clusters were determined at a resolution of 0.5.Seurat’s *FindAllMarkers* function was used to perform differential gene expression (DEG) analysis and compare each cluster against all other clusters by leveraging the Wilcoxon rank sum test. Marker genes were identified using a stringent filtering criterion, including minimum percentage expression of 0.25, a log fold change threshold of 0.25 and an adjusted p-value of 0.05. The top 50 marker genes significantly enriched were used to infer potential cell types and annotate individual clusters using EnrichR and the PanglaoDB gene database (Chen et al., 2013; Kuleshov et al., 2016; Franzen et al ., 2019; Xie et al., 2021).

### Pseudo-bulk Processing and Differential Gene Expression

To perform a more comprehensive DEG analysis between clusters, gene expression levels from individual spots within each cluster were aggregated. Using the *scater* package, spots were pseudo-bulked by summing raw UMI counts of each gene within each cluster (McCarthy et al., 2017). To increase statistical power, the clusters were pooled into ten pseudo-replicates. Using *filterByExpr* and *calcNormFactors*, implemented within the *edgeR* package, genes with insufficiently low counts in the pseudo-bulked clusters were removed and read counts were normalised (Chen et al., 2024). Subsequently, DEG analysis was conducted using the *edgeR* framework, incorporating elements of linear modelling from the *limma* package for robust statistical inference, allowing genes with significantly altered expression levels between each cluster to be identified (Ritchie et al. 2015). *edgeR* further fits a negative binomial model using *glmQLFit* to account for overdispersion and employs *glmTreat* to identify differentially expressed genes based on a false discovery rate (FDR)-adjusted p-value < 0.05 and a logFC threshold of 1.2. All upregulated marker genes were included in the analysis, except for those in cluster 5, where only the top 50 genes ranked by FDR were considered. To interpret results, a heatmap was generated using *pheatmap*, providing a visual representation of differentially expressed gene expression patterns across each cluster.

### Cell Cycle Analysis

Cell cycle phase assessment was performed using Seurat’s *CellCycleScoring* function. Pre-defined gene sets specific to human were employed to identify cells in S and G2/M phases as per the scoring strategy described by Tirosh et al, (2016).

### Immune Cell Type Identification

Based on previous literature, manually defined marker gene sets were used to determine significantly different expression patterns and perform gene set enrichment analysis (GSEA) for each cluster. Immune cell populations were identified by the following markers: T-regs (*FOXP3*, *TGFb* and *IL10*) (Yano et al., 2019; Enk, A., 2005; Sun et al., 2023), anti-inflammatory T cells (*IFNg*, *IL4*, *IL5* and *IL13*) (Al-Quahtrani et al., 2024; Sun et al., 2023), pro-inflammatory T cells (*IL10*) (Al-Quahtrani et al., 2024; Sun et al., 2023), M1 macrophages (*CD86*, *IL1b*, *IL6*, *TNFa*, *INFy*, *CCL2*, *CCL9* and *CCL10*) (Al-Quahtrani et al., 2024; Colombo et al., 2024; Sun et al., 2023), M2 macrophages (*IL10*, *TGFb*, *IL4*, *IL13*, *CCL17* and *CCL22*) (Al-Quahtrani et al., 2024; Colombo et al., 2024; Sun et al., 2023) and Langerhans cells (*CD207*, *CD1a*) (Maarifi et al., 2020; Mizumoto & Takashima, 2004). Using *AddModuleScore*, co-expression patterns of gene set markers are identified and adjusted by subtracting the aggregated expression across the whole tissue sample (Tirosh et al., 2016).

### Macrophage Activity Score

Macrophage activity within cluster 8, 9 and 10 were calculated on the predefined marker gene list for M1 macrophages and M2 macrophages, as listed above. The difference in macrophage activity between the two cell states within these clusters were visualised using a boxplot (Figure 2d.).

**Figure 1.**
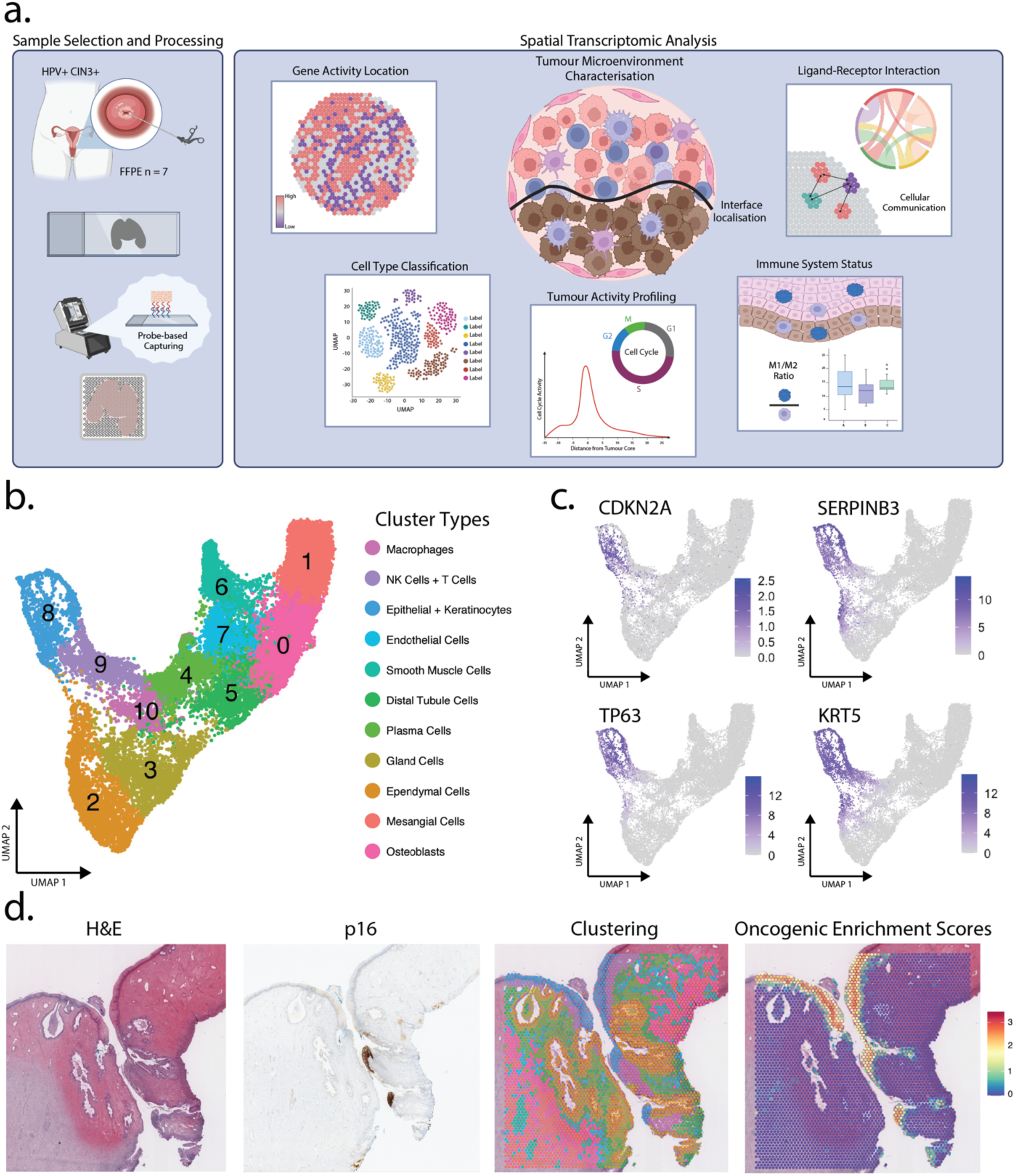
Characterising the immuno-spatial landscape of seven HPV+ CIN3+ samples. a) Schematic of experimental design and spatial transcriptomic analysis. b) Integrated UMAP displaying cell type distribution across the seven samples. c) Feature plots of known cervical cancer oncogenes: *CDKN2A*, *SERPINB3*, *TP63* and *KRT5* (Ou et al., 2021). d) A representative sample (VLP89_A) with H&E image, p16 immunohistochemistry staining for CIN3+ lesion validation, spatial clustering (with corresponding cluster colours as clusters shown in panel b) and identification of CIN3+ region using cervical cancer oncogene co-expression.

**Figure 2.**
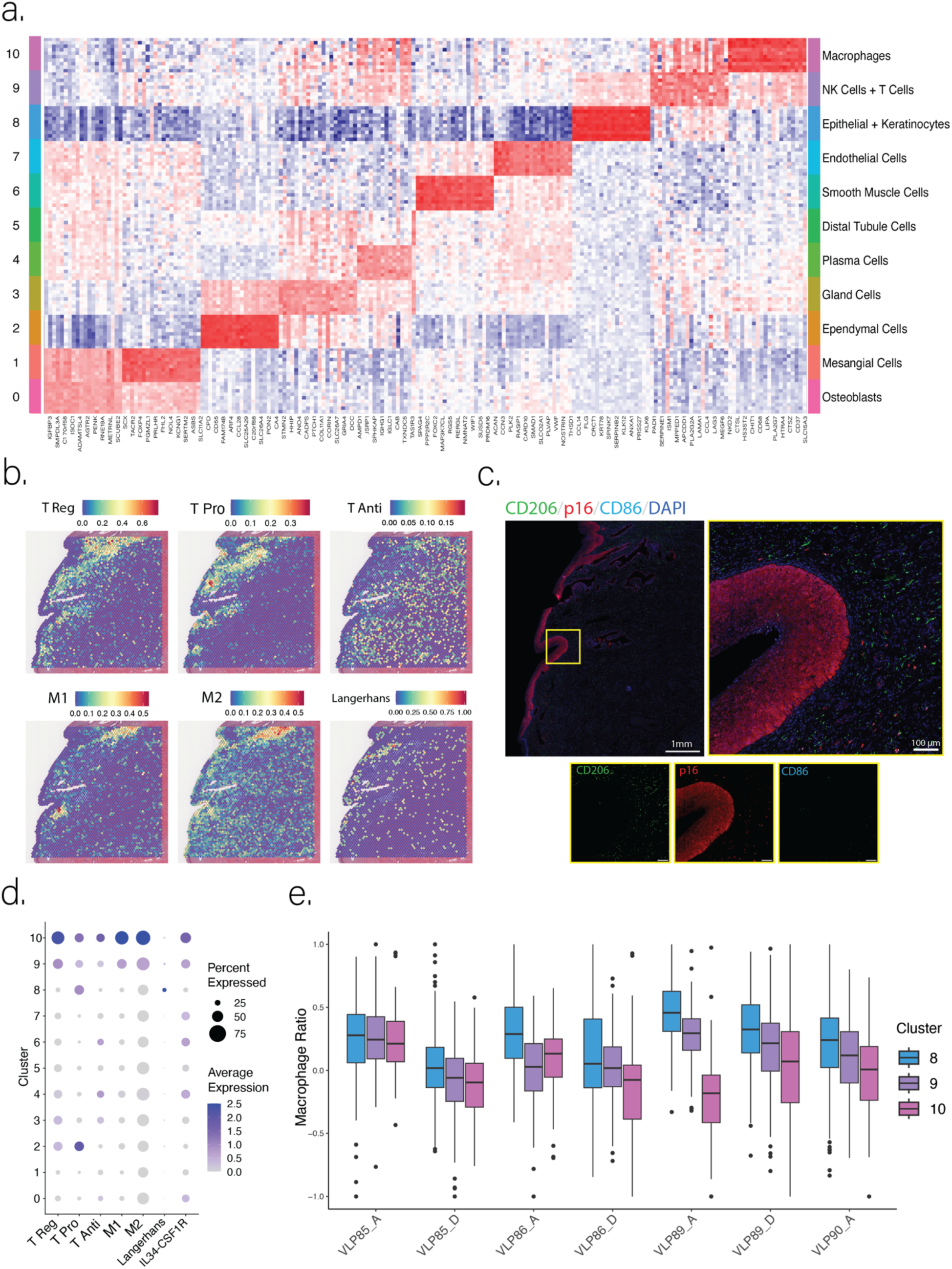
Analysis of immune cell types and identification of a distinct immune cluster. a) Expression heatmap of top 10 upregulated marker genes for each cluster. b) Spatial mapping of immune cell module scores for a representative sample (VLP86_A). c) Immunofluorescent staining on adjacent tissue sections for representative sample (VLP86_A) to evaluate the CIN3+ lesion (p16) and presence of M1 (CD86) and M2 (CD206) macrophages. Nuclei were stained by DAPI. Fluorescent signals for a magnified region, highlighted by the yellow box, shown in grayscale for each channel with scale bars indicating 100 µm. d) Dot plot of gene expression scores based on canonical immune cell markers. e) M2 to M1 module score ratio in clusters 8, 9 and 10 for all seven samples.

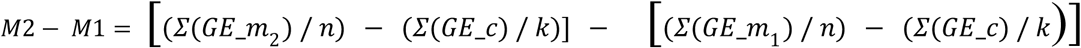

*GE_m_1_*: Expression value of genes in M1 macrophage module group

*GE_m_2_*: Expression value of genes in M2 macrophage module group

*GE_m_c_*: Expression value of genes in the control group for module

*n*: Number of genes in each module group

*k*: Number of genes in the control group

### IL34-CSF1R vs Tumour Expression

The relationship between IL34-CSF1R co-expression and the tumour microenvironment was investigated by comparing the spatial expression patterns of IL34-CSF1R and cervical cancer gene markers (*SERPINB3*, *TP63*, *KRT5* and *CDKN2A*) (Ou et al., 2021). To determine meaningful expression, only data points with a module score above a threshold of 0.1 were considered, otherwise the spots were labelled as “None”. Subsequently, through a module-based data partitioning method, remaining spots were categorised into “IL34-CSF1R” or “Tumour” based on their dominant module score.

### Pathway Analysis

For each cluster, we used ClusterProfiler to identify Gene Ontology (GO) terms related to biological processes that were significantly enriched related to the respective DEG lists. The *enrichGO* function and the human genome database, were employed to determine significantly enriched terms with a p-value lower than 0.01. The most significant GO term was assigned to each cluster and compared it to the existing cluster annotation label. The identified terms were categorised into common ontology tags and visualised using *REVIGO* (Supek et al., 2011).

### Cell-Cell Interaction Analysis

Using stLearn, a package designed for downstream ST analysis, cellular communication is inferred through ligand-receptor interactions within and between spots (Pham et al., 2023). From a list of known ligand-receptor pairs, the package calculates the genetic expression for each ligand and receptor and determines the statistical difference of the ligand-receptor co-expression. In comparison to the background distribution of all random gene-gene pairs, the p-value is determined by the cumulative occurrence where the ligand-receptor pair has scored higher than other random combination pairs. The spatial proximity of each ligand-receptor pair is also taken into consideration, with intermediate neighbouring cells more likely to interact. By combining genetic expression and spatial information, stLearn is able to rank and assess the significance of ligand-receptor pair interactions and interpret potential cell-cell communication.

### Oropharyngeal Squamous Cell Carcinoma (OPSCC) Analysis

IL34-CSF1R expression was further analysed in a public oropharyngeal squamous cell carcinoma (OPSCC) dataset to explore potential similarities in HPV+ carcinogenesis (Causer et al., 2023). A module score for the expression of OPSCC HPV+ tumour marker genes (*SOX4, SLC2A1, KRT8, EIF4A2, SNAI2, CA12, HSPH1, ANGPTL4, FAM162A, PFKP, TGFB1, UHRF2, PSIP1, MYC, RCL1, KDM4C, ACTL6A, CHAF1A, DEK, ECT2, POLR1D, GAS1, PLOD2, DDIT4, IL33, ABCC5, ALCAM, CALCRL, MKNK2, MT2A*) and IL34-CSF1R co-expression was determined and plotted using *SpatialDimPlot*. Gene set expression was compared with pathological annotations associated with OPSCC HPV+ samples (Causer et al., 2023).

## Results

### Unsupervised identification of 11 gene expression clusters across seven CIN3+ patients

Utilising the comprehensive transcriptome-wide profiling capabilities of Visium CytAssist (10X Genomics), the spatially resolved gene expression profiles of CIN3+ lesions were captured using a probe-based approach (Figure 1a). Across the seven samples, the transcriptome for 21,054 barcoded 55µm diameter spots were measured, with each spot containing approximately one to ten cells. A median of 4,422 – 5,730 genes were detected per spot, demonstrating transcriptionally active cells across the tissue. Unsupervised clustering revealed 11 distinct clusters amongst seven samples which were then annotated based on gene set enrichment analysis using the PanglaoDB cell type database (Figure 1b), (Franzen et al., 2019). The identified clusters were annotated with major cell types as osteoblasts (0), mesangial cells (1), ependymal cells (2), gland cells (3), plasma cells (4), distal tubule cells (5), smooth muscle cells (6), endothelial cells (7), epithelial cells and keratinocytes (8), natural killer cells and T cells (9), and macrophages (10) (Figure 2a). Although inter-patient heterogeneity was evident in the variations in the distribution of cell types between samples, all samples exhibited similar patterns in cluster formation and gene expression (Sup Fig 1). To validate the clustering results, we employed both gene set enrichment analysis and gene ontology databases, confirming the accuracy of cell type identification (Figure 2a).

### Mapping the neoplastic cluster highly enriched for oncogenes

We observed that cluster 8, representing keratinocytes, contained the highest number of significant differentially expressed genes when compared to the other clusters, indicating a unique microenvironment with distinct cell activity (Figure 1a). Genes most up-regulated by cluster 8 included *KRT78, SERPINB2*, *ANXA1* and *CLIC3*. Additional differentially expressed genes that were over-expressed in cluster 8 included known canonical cervical cancer oncogenes (Ou et al., 2021): *CDKN2A*, *SERPINB3*, *TP63* and *KRT5* (Figure 1c). Spatial expression patterns of *SERPINB3*, *TP63*, and *KRT5* showed particularly high co-localisation with cluster 8, suggesting the presence of neoplastic epithelial cells within this region (Figure 1d and Sup Fig 1 for all seven samples). However, the spatial distribution of *CDKN2A,* encoding tumour suppressor proteins p16INK4a and p14arf, displayed greater heterogeneity between each sample (Sup Fig 2). Increased LC-specific gene expression was also observed in the neoplastic cluster 8 region in comparison to other clusters (Figure 2b). Cell cycle scoring demonstrated high proportions of spots in S phase within this defined neoplastic region, indicating active cell replication (refer to Sup Fig 1 for the location of cluster 8 represented by the blue-coloured spots, Sup Fig 3).

### Unique community displaying increased immune cell activity

Gene expression analysis revealed prominent immune cell activity within cluster 10, based on canonical immune marker genes (Sup Table 1). Based on differentially expressed genes, macrophages were observed as the dominant cell type (Figure 2a). In addition, T cell associated genes (Sup Table 1) were enriched within cluster 10 in comparison to other clusters (Figure 2d). The expression signature for the main immune cell types, (i.e., scores based on multiple immune markers – refer to the method session), is shown in Figure 2b. Spatial profiling demonstrated that the immune region (cluster 10) regularly neighboured the neoplastic region (cluster 8) across the seven samples (Sup Fig 1). Further sub-clustering of cluster 10 differentiated this cluster into spots with M1 and M2 macrophage gene signatures. Cell type proportions of these macrophages and other immune cell types displayed heterogeneity between samples (Sup Fig 4). Using relative geneset enrichment scores, we calculated M2/M1 macrophage ratios for each spot across individual tissue sections. The M2/M1 ratio was consistently higher in the neoplastic region (cluster 8) across all samples, indicating an immune-suppressive macrophage microenvironment located specifically to the neoplasia (Figure 2e). This differential macrophage signature was corroborated by immunofluorescence staining of CD206 (M2) and CD86 (M1) in adjacent tissue sections, where M2 macrophages were more abundant around the CIN3 lesions (positive for p16) compared to M1 macrophages (Figure 2c). Inversely, the adjacent immune regions within cluster 10 had a comparatively low M2/M1 ratio across all samples, suggesting a more pro-inflammatory M1 macrophage microenvironment localised outside of the neoplastic region (Figure 2e). Cluster 9 also demonstrated high immune cell activity, with an intermediate M2/M1 ratio between that of the neoplastic region (clusters 8) and the immune region (cluster 10) (Figure 2e). The immune activity scores across all samples were evaluated in Sup Fig 5. Cluster 9 also consistently bordered the neoplastic region and, at times, formed a complete barrier between cluster 8 and 10 when inspected spatially, highlighting a possible transitional site between normal and neoplastic regions (Figure 1d).

### Cell-cell interaction and ligand-receptor pair analysis

We performed transcriptome-wide cell-cell interaction (CCI) analysis to reveal interaction hot spots and consistent interactions between ligand-receptor pairs and cell type pairs across all samples. Cluster 10, enriched for immune cells, displayed the lowest cell-type frequency across all samples (Figure 3a), but demonstrated the highest level of ligand-receptor activity, predominantly intra-cluster interactions (Figure 3b). Interestingly, the neoplastic cluster 8 demonstrated lower CCI activity (Figure 3a), with most interactions occurring between either cluster 9 or with itself (Figure 3b). Notably, both cluster 8 and 10 demonstrated high inter-cluster relationships with cluster 9, but almost no interactions occurred directly between cluster 8 and 10 (Figure 3b). Specifically, 61 ligand-receptor interactions were observed between clusters 8 and 9, 75 interactions between clusters 9 and 10, and only 4 interactions between clusters 8 and 10 (Sup Fig 6). The top molecule signalling between neoplastic cluster 8 and the adjacent cluster 9 was Midkine (MDK), a well-recognised gene overexpressed by various human malignancies including cervical cancer and playing a role in cell growth, survival metastasis, migration and angiogenesis (Moon et al., 2003). The S100A9-TLR4 signalling axis has been implicated in inducing immunosuppressive myeloid-derived suppressor cells (Özbay Kurt et al., 2024), and TRAIL signalling, while being a death signal, has been implicated in cell migration and invasion thereby promoting cancer progression (Peyre et al., 2021). The identified communication between the neoplastic region (cluster 8) and the immune region (cluster 10) suggests that the viral presence keeps inducing a pro-inflammatory response in form of complement activity, type 1 interferon signalling and XCL2-mediated immune cell recruitment, while APOE signalling favours immune suppression (Sup Fig. 6). Overall, this data highlights the complex molecular and spatial intricacies in immune recognition of the virally-induced abnormalities of the neoplastic region, and the signalling from this neoplastic region to suppress immune activity and promote its progression towards an invasive phenotype.

**Figure 3.**
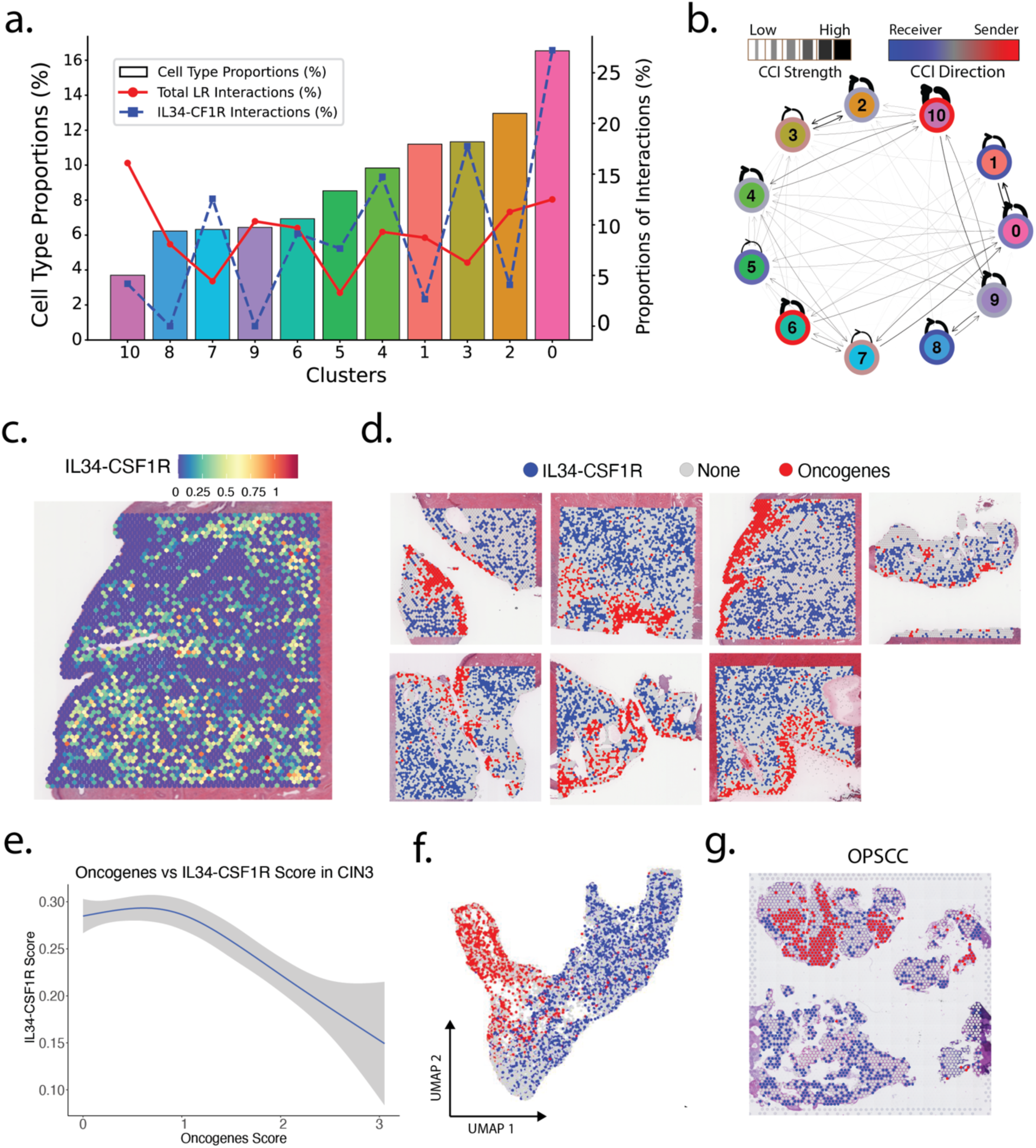
Spatial analysis of cell-cell interactions based on neighbourhood expression of ligand-receptor pairs in CIN3+ samples and comparisons with external OPSCC samples. **a)** Cell type proportions (bar plots) for each cluster across all samples with the percentage of ligand-receptor interactions (line plots). The total ligand-receptor interactions are shown as the red line, (proportion of spots with significant interactions among all spots) and the IL34-CSF1R interactions as shown as the blue line (significant IL34-CSF1R interactions among significant interactions of all ligand-receptor pairs). **b)** Cell-cell interaction network displaying intra- and inter-cluster ligand-receptor interactions with thicker lines representing stronger interactions and arrows indicate receiver/sender status of each cluster pair. **c)** Spatial visualisation of IL34-CSF1R co-expression module score (refer to Methods). **d)** Cancer vs IL34-CSF1R dominating region categorisation determined by preponderant module score expression of either cancer oncogenes, IL34-CSF1R co-expression or neither within each spot for all seven samples. **e)** Changes in IL34-CSF1R interaction scores relative to cervical cancer module scores. **f)** Integrated UMAP displaying distribution of spots categorised by higher co-expression tumour oncogenes (red) or IL34-CSF1R across all samples (blue). **g)** Comparison of cancer oncogenes vs Il34-CSF1R spatial expression patterns in a different HPV+ cancer system, namely OPSCC tissues with the same categorical colourisation as panel d). The OPSCC data was generated by Causer et al. (2023).

### Spatial gene expression patterns of IL34 and CSF1R suggest their interactions and roles in tumour progression

Our previous research has identified IL34 as a dysregulated and prognosis-associated gene in CIN and cervical cancer (Tuong et al., 2021). We therefore aimed to define the spatial characteristics of IL34 signalling in this dataset. The IL34-CSF1R ligand-receptor relationship was explored within clusters by comparing co-expression levels respective to cancer oncogene co-expression. The highest IL34-CSF1R interaction activity was observed in cluster 0, followed by cluster 3 (Figure 3a). Notably, we observed the lowest co-expression of this ligand-receptor pair within the neoplastic and adjacent transitional regions of cluster 8 and cluster 9 (Figure 3a). When plotted spatially, the co-expression of Il34-CSF1R was seemingly absent in neoplastic regions (Figure 3c), where the expression of cancer oncogenes dominated (Figure 3d). A negative correlation was observed between co-expression of IL34-CSF1R and cancer oncogenes (Figure 3e) with a separation of gene expression when visualised on the UMAP (Figure 3f). Importantly though, our data suggests that the IL34-receiving cells (LCs and macrophages) are not absent from these neoplastic regions that lack IL34 (Fig. 2b, c, d), indicating that other molecules may contribute to LC retainment, macrophage infiltration and M2 polarisation. To see whether the IL34-CSF1R interaction pattern extended to other HPV-mediated cancer type, we explored a publicly available oropharyngeal cancer sample (Causer et al., 2023), which demonstrated similarities in regard to the absence of IL34-CSF1R co-expression in neoplastic regions (Figure 3g). This data is in line with our previous research demonstrating a loss of IL34 in CIN, cervical cancer (Tuong et al., 2021), in head and neck squamous cell cancers and a murine model of HPV-mediated epithelial hyperplasia (unpublished Dang et al., n.d.), and spatially defines the strict refinement of IL34 loss to the neoplastic regions while maintaining signalling in adjacent tissue regions.

## Discussion

Although cervical cancer incidence has decreased over the past decades, it remains a significant global burden, with an estimated 604,127 new cases and 341,831 deaths worldwide in 2020 (Singh et al., 2023). Despite the emergence of medical advancements, there remains ambiguity surrounding the mechanisms underlying HPV persistence and disease progression. We set out to conduct a detailed spatial analysis of human CIN3 samples, focusing on the complex interplay between immune cells and the tumour microenvironment, particularly the role of IL34 and CSF1R in tumour progression as proposed in previous research.

We defined the presence of a neoplastic region across our dataset, cluster 8, characterized by differentially expressed genes, including known cervical cancer oncogenes. Cluster 8 was most enriched for *KRT78* and *SERPINB2,* both of which have been shown to exhibit increased expression in HPV-associated cervical cancers (Varga et al., 2017; Syrjänen et al., 2009). Notably, elevated *SERPINB2* expression has been linked to the severity of cervical lesions (Syrjänen et al., 2009). Furthermore, we observed an upregulation of *ANXA1*, which has been shown to modulate inflammatory properties by promoting macrophage differentiation to its pro-tumoral M2 phenotype, promoting tumour progression (Araújo et al., 2021; Moraes et al., 2017). Additionally, *CLIC3* was significantly upregulated in this cluster, a gene associated with HPV-induced neoplastic cervical lesions and contributes to poor prognosis in patients with cervical carcinoma (Guan et al., 2019; Steenbergen et al., 2013). We also observed the enrichment of *CDKN2A* (*p16*), *SERPINB3*, *KRT5* and *TP63*, which are well-established biomarkers for HPV-positive neoplasia and are typically expressed in tumour areas (Ou et al., 2021). KRT5 and TP63 are known epithelial and basal markers, and their overexpression has been implicated in pathways and lineages that drive carcinoma progression (Chumduri et al., 2021; Jiang et al., 2024). CDKN2A is a canonical proliferation marker that is also a well-documented biomarker for HPV infection, with expression levels increasing from early to late-stage CIN and furthermore to invasive cervical cancer (Chernock et al., 2013; Liu et al., 2023, Wu & Xi, 2021). Additionally, SERPINB3 has recently been associated with poor prognosis in cervical cancer (Fan et al., 2023). SERPINB3 is believed to contribute to tumour progression by protecting cancer cells from immune surveillance, thereby facilitating therapeutic evasion. The elevated expression of SERPINB3 within cluster 8, the neoplastic region, suggests a potential mechanism for immune escape and treatment resistance in this subset of cervical cancer cells. Cell-cycle analysis indicated that this determined neoplastic cluster was predominantly in the ‘S’ or DNA replication phase, consistent with the proliferative nature of CIN3 lesions (Pinto et al., 2008; Singh et al., 2008).

We observed an immune cell cluster (cluster 10) consistently neighbouring the lesions in each sample. Cluster 10 exhibited heightened immune cell activity, particularly T cells, macrophages, and LCs, suggesting an active and localised immune activity. Further analysis showed that a M2 macrophage cell state was more prevalent in the neoplastic region compared to normal adjacent regions, aligning with previous investigations exploring the spatiotemporal expression of M1/M2 in cervical cancer tissues (Guo et al., 2023). This finding supports previous studies highlighting the immunosuppressive role of M2 macrophages in promoting tumour growth and progression (Guo et al., 2023; Tuong et al., 2021).

Our study identified several ligand-receptor pairs that may play crucial roles in cervical tumorigenesis and immune modulation. Ligand-receptor analysis revealed a high number of cell-cell interactions in the immune cell cluster 10 but minimal interactions between the neoplastic cluster 8 and the immune cell cluster 10. For instance, the MDK-SDC1 interaction, observed in our samples between the neoplastic cluster 8 and the adjacent cluster 9, has previously been reported as a prognostic biomarker (Oto et al., 2023). MDK-TSPAN1 has been shown to promote tumorigenesis and chemoresistance in head and neck cancer (Huang et al., 2008; Garcia-Meyea et al., 2020). The statistically significant interactions using these ligand-receptor pairs in CIN3 lesions suggests their potential involvement in cervical cancer development and immune evasion.

The distinct coexpression patterns of IL34 and CSF1R suggest their involvement in tumour progression, immune suppression, and potentially macrophage polarisation. Although IL34 was previously thought to be solely a secreted protein because it lacks transmembrane domains in its primary structure (Lin et al., 2008), Ogawa et al. discovered a membrane-bound form of IL34 through anti-mouse IL34 antibody staining. It was found that only the cell-surface IL34, and not the secreted variant, binds to the CSF1R receptor and induces the maturation of follicular dendritic cell-induced monocytic cells (Ogawa et al., 2019). This implies that the membrane-bound form is the active form of IL34, and cell-to-cell contact is required to convey function. We observed a lack of co-expression of IL34 and CSF1R within the neoplastic regions, despite high expression in the adjacent tissues. Additionally, there was an overall abundance of CSF1R compared to IL34, suggesting that macrophage infiltration in the neoplastic region occurs despite the absence of IL34. However, neoplastic regions absent of IL34 were enriched primarily with M2 gene signatures, while adjacent regions rich in IL34 expression were enriched with M1 gene signature, suggesting that the lack of IL34 may result in accumulation of immune-suppressive macrophage phenotypes. This contrasts with the expression pattern of this ligand-receptor pair observed in melanoma, where it has been described that the heightened expression of IL34 is related to lower immune cell frequencies, suggesting a more protumorigenic role (Alshaebi et al., 2023; Han et al., 2018, unpublished Grice et al., n.d.). This suggests a unique mechanism of action for HPV-driven carcinogenesis. To assess the generalisation of this finding, we also conducted additional analysis and observed parallels in IL34-CSF1R expression between OPSCC and CIN3, suggesting similarities in HPV+ driven neoplasia regardless of the anatomical site (Causer et al., 2023). This highlights the potential for translational research across different types of squamous cell carcinomas.

Our study has several limitations, including a small sample size, and the lack of different CIN stages. These limitations may affect the generalizability of our findings and the interpretation of spatial and gene expression patterns. Future studies with larger cohorts, encompassing disease progression from normal cervical to cervical cancer and incorporating more diverse omics analyses are needed to validate and extend upon our observations. Despite the heterogeneity, our deep analysis of spatial transcriptomics data across the seven human CIN3 samples reveals consistent spatial patterns of the immune signatures within and adjacent to the precancer microenvironment. We found distinct and spatially defined interaction activities relative to locations with an active HPV+ signature. Together, the results suggest a potential strategy to modulate the immune system in HPV-positive patients to prevent progression. This promising approach warrants further investigation.

## Acknowledgement

We thank Dr. Diane Cominos at Sullivan Nicolaides Pathology and Prof. Andreas Obermair at Queensland Centre for Gynaecological Cancer (QCGC) for providing samples and supports in this project.

## References

1. Abd Warif, N. M., Stoitzner, P., Leggatt, G. R., Mattarollo, S. R., Frazer, I. H., & Hibma, M. H. (2015). Langerhans cell homeostasis and activation is altered in hyperplastic human papillomavirus type 16 E7 expressing epidermis. PloS one, 10(5), e0127155. 10.1371/journal.pone.0127155

2. Al-Qahtani, A. A., Alhamlan, F. S., & Al-Qahtani, A. A. (2024). Pro-inflammatory and anti-inflammatory interleukins in infectious diseases: A comprehensive review. Tropical Medicine and Infectious Disease, 9(1), 13. 10.3390/tropicalmed9010013

3. Alshaebi, F., Safi, M., Algabri, Y. A., Al-Azab, M., Aldanakh, A., Alradhi, M., Reem, A., & Zhang, C. (2023). Interleukin-34 and immune checkpoint inhibitors: Unified weapons against cancer. *Frontiers in Oncology, 13*, 1099696. 10.3389/fonc.2023.1099696

4. Araújo, T. G., Mota, S. T. S., Ferreira, H. S. V., Ribeiro, M. A., Goulart, L. R., & Vecchi, L. (2021). Annexin A1 as a Regulator of Immune Response in Cancer. Cells, 10(9), 2245. 10.3390/cells10092245

5. Baghdadi, M., Umeyama, Y., Hama, N., Kobayashi, T., Han, N., Wada, H., & Seino, K. I. (2018). Interleukin-34, a comprehensive review. Journal of leukocyte biology, 104(5), 931–951. 10.1002/JLB.MR1117-457R

6. Bashaw, A. A., Teoh, S. M., Tuong, Z. K., Leggatt, G. R., Frazer, I. H., & Chandra, J. (2019). HPV16 E7-Driven Epithelial Hyperplasia Promotes Impaired Antigen Presentation and Regulatory T-Cell Development. The Journal of investigative dermatology, 139(12), 2467– 2476.e3. 10.1016/j.jid.2019.03.1162

7. Bashaw, A. A., Zhou, C., Yu, M., Tolley, L., Leggatt, G. R., Frazer, I. H., & Chandra, J. (2021). Regulatory T Cells but Not IL-10 Impair Cell-Mediated Immunity in Human Papillomavirus E7+ Hyperplastic Epithelium. The Journal of investigative dermatology, 141(5), 1264–1273.e3. 10.1016/j.jid.2020.10.011

8. Brisson, M., & Drolet, M. (2019). Global elimination of cervical cancer as a public health problem. *The Lancet Oncology, 20*(3), 319–321. 10.1016/S1470-2045(19)30072-5

9. Cattani, P., Zannoni, G. F., Ricci, C., D’Onghia, S., Trivellizzi, I. N., Di Franco, A., Vellone, V. G., Durante, M., Fadda, G., Scambia, G., Capelli, G., & De Vincenzo, R. (2009). Clinical performance of human papillomavirus E6 and E7 mRNA testing for high-grade lesions of the cervix. *Journal of Clinical Microbiology, 47*(12), 3895–3901. 10.1128/JCM.01275-09

10. Cersosimo, F., Lonardi, S., Ulivieri, C., Martini, P., Morrione, A., Vermi, W., Giordano, A., & Giurisato, E. (2024). CSF-1R in Cancer: More than a Myeloid Cell Receptor. Cancers, 16(2), 282. 10.3390/cancers16020282

11. Chandra, J., Miao, Y., Romoff, N., & Frazer, I. H. (2016). Epithelium Expressing the E7 Oncoprotein of HPV16 Attracts Immune-Modulatory Dendritic Cells to the Skin and Suppresses Their Antigen-Processing Capacity. PloS one, 11(3), e0152886. 10.1371/journal.pone.0152886

12. Chen, E. Y., Tan, C. M., Kou, Y., Duan, Q., Wang, Z., Meirelles, G. V., Clark, N. R., & Ma’ayan, A. (2013). Enrichr: Interactive and collaborative HTML5 gene list enrichment analysis tool. *BMC Bioinformatics, 14*, 128. 10.1186/1471-2105-14-128

13. Chen, L., Shi, V., Wang, S., Sun, L., Freeman, R., Yang, J., Inkman, M. J., Ghosh, S., Ruiz, F., Jayachandran, K., Huang, Y., Luo, J., Zhang, J., Cosper, P., Luke, C. J., Spina, C. S., Grigsby, P. W., Schwarz, J. K., & Markovina, S. (2023). SCCA1/SERPINB3 suppresses antitumor immunity and blunts therapy-induced T cell responses via STAT-dependent chemokine production. The Journal of clinical investigation, 133(15), e163841. 10.1172/JCI163841

14. Chen, Y., Chen, L., Lun, A. T. L., Baldoni, P., & Smyth, G. K. (2024). edgeR 4.0: Powerful differential analysis of sequencing data with expanded functionality and improved support for small counts and larger datasets. *bioRxiv*. 10.1101/2024.01.21.576131

15. Chernock, R. D., Wang, X., Gao, G., Lewis, J. S., Jr, Zhang, Q., Thorstad, W. L., & El-Mofty, S. K. (2013). Detection and significance of human papillomavirus, CDKN2A(p16) and CDKN1A(p21) expression in squamous cell carcinoma of the larynx. Modern pathology : an official journal of the United States and Canadian Academy of Pathology, Inc, 26(2), 223–231. 10.1038/modpathol.2012.159

16. Chumduri, C., Gurumurthy, R. K., Berger, H., Dietrich, O., Kumar, N., Koster, S., Brinkmann, V., Hoffmann, K., Drabkina, M., Arampatzi, P., Son, D., Klemm, U., Mollenkopf, H. J., Herbst, H., Mangler, M., Vogel, J., Saliba, A. E., & Meyer, T. F. (2021). Opposing Wnt signals regulate cervical squamocolumnar homeostasis and emergence of metaplasia. Nature cell biology, 23(2), 184–197. 10.1038/s41556-020-00619-0

17. Colombo, G., Pessolano, E., Talmon, M., Genazzani, A. A., & Kunderfranco, P. (2024). Getting everyone to agree on gene signatures for murine macrophage polarization in vitro. PloS one, 19(2), e0297872. 10.1371/journal.pone.0297872

18. Enk, A. (2005). Dendritic cells in tolerance induction. Immunology Letters 99(1) 8–11. 10.1016/j.imlet.2005.01.011

19. Franzén O, Gan LM, Björkegren JLM. (2019). PanglaoDB: a web server for exploration of mouse and human single-cell RNA sequencing data. Database (Oxford). 1; 2019:baz046. doi: 10.1093/database/baz046.

20. Garcia-Mayea, Y., Mir, C., Carballo, L., Castellvi, J., Temprana-Salvador, J., Lorente, J., Benavente, S., García-Pedrero, J. M., Allonca, E., Rodrigo, J. P., & LLeonart, M. E. (2020). TSPAN1: A novel protein involved in head and neck squamous cell carcinoma chemoresistance. *Cancers, 12*(11), 3269. 10.3390/cancers12113269

21. Greter, M., Lelios, I., Pelczar, P., Hoeffel, G., Price, J., Leboeuf, M., Kündig, T. M., Frei, K., Ginhoux, F., Merad, M., & Becher, B. (2012). Stroma-derived interleukin-34 controls the development and maintenance of langerhans cells and the maintenance of microglia. Immunity, 37(6), 1050–1060. 10.1016/j.immuni.2012.11.001

22. Guan, Y. T., Xie, Y., Zhou, H., Shi, H. Y., Zhu, Y. Y., Zhang, X. L., Luan, Y., Shen, X. M., Chen, Y. P., Xu, L. J., Lin, Z. Q., & Wang, G. (2019). Overexpression of chloride channel-3 (ClC-3) is associated with human cervical carcinoma development and prognosis. Cancer cell international, 19, 8. 10.1186/s12935-018-0721-x

23. Guo, C., Qu, X., Tang, X., Song, Y., Wang, J., Hua, K., & Qiu, J. (2023). Spatiotemporally deciphering the mysterious mechanism of persistent HPV-induced malignant transition and immune remodelling from HPV-infected normal cervix, precancer to cervical cancer: Integrating single-cell RNA-sequencing and spatial transcriptome. *Clinical and Translational Medicine, 13*(3), e1219. 10.1002/ctm2.1219

24. Hafemeister, C., & Satija, R. (2019). Normalization and variance stabilization of single-cell RNA-seq data using regularized negative binomial regression. *Genome Biology, 20*, 296. 10.1186/s13059-019-1874-1

25. Han, N., Baghdadi, M., Ishikawa, K., Endo, H., Kobayashi, T., Wada, H., Imafuku, K., & Seino, K. I. (2018). Enhanced IL-34 expression in Nivolumab-resistant metastatic melanoma. *Inflammation and Regeneration, 38*, 3. 10.1186/s41232-018-0060-2

26. Hockey, L., Mulay, O., Xiong, Z., Tan, X. S., Khosrotehrani, K., Nefzger, M. C. & Nguyen, Q. (2024) MMCCI: Multimodal Integrative Analysis of single-cell and spatial cell-type communications to uncover overarching and condition-specific ligand-receptor interaction pathways [Preprint]. doi:10.1101/2024.02.28.582639.

27. Huang, Y., Sook-Kim, M., & Ratovitski, E. (2008). Midkine promotes tetraspanin-integrin interaction and induces FAK-Stat1alpha pathway contributing to migration/invasiveness of human head and neck squamous cell carcinoma cells. *Biochemical and Biophysical Research Communications, 377*(2), 474–478. 10.1016/j.bbrc.2008.09.138

28. Hume, D. A., Gutowska-Ding, M. W., Garcia-Morales, C., Kebede, A., Bamidele, O., Vallejo Trujillo, A., & Smith, J. (2020). Functional evolution of the colony-stimulating factor 1 receptor (CSF1R) and its ligands in birds. Journal of Leukocyte Biology, 107(2), 237–250. 10.1002/JLB.6MA0519-172R

29. Itay, T., et al. (2016). Dissecting the multicellular ecosystem of metastatic melanoma by single-cell RNA-seq. *Science, 352*, 189-196. 10.1126/science.aad0501

30. Jiang, Y., Zheng, Y., Zhang, Y. W., Kong, S., Dong, J., Wang, F., Ziman, B., Gery, S., Hao, J. J., Zhou, D., Zhou, J., Ho, A. S., Sinha, U. K., Chen, J., Zhang, S., Yin, C., Wei, D. D., Hazawa, M., Pan, H., Lu, Z., … Jiang, Y. Y. (2024). Reciprocal inhibition between TP63 and STAT1 regulates anti-tumor immune response through interferon-γ signaling in squamous cancer. Nature communications, 15(1), 2484. 10.1038/s41467-024-46785-9

31. Kuleshov, M. V., Jones, M. R., Rouillard, A. D., Fernandez, N. F., Duan, Q., Wang, Z., Koplev, S., Jenkins, S. L., Jagodnik, K. M., Lachmann, A., McDermott, M. G., Monteiro, C. D., Gundersen, G. W., & Ma’ayan, A. (2016). Enrichr: A comprehensive gene set enrichment analysis web server 2016 update. *Nucleic Acids Research, 44*(W1), W90-W97. 10.1093/nar/gkw377

32. Kurmyshkina, O. V., Dobrynin, P. V., Kovchur, P. I., & Volkova, T. O. (2023). Sequencing-based transcriptome analysis reveals diversification of immune response- and angiogenesis-related expression patterns of early-stage cervical carcinoma as compared with high-grade CIN. *Frontiers in Immunology, 14*, 1215607. 10.3389/fimmu.2023.1215607

33. Lelios, I., Cansever, D., Utz, S. G., Mildenberger, W., Stifter, S. A., & Greter, M. (2020). Emerging roles of IL-34 in health and disease. The Journal of experimental medicine, 217(3), e20190290. 10.1084/jem.20190290

34. Lin, H., Lee, E., Hestir, K., Leo, C., Huang, M., Bosch, E., Halenbeck, R., Wu, G., Zhou, A., Behrens, D., Hollenbaugh, D., Linnemann, T., Qin, M., Wong, J., Chu, K., Doberstein, S. K., & Williams, L. T. (2008). Discovery of a cytokine and its receptor by functional screening of the extracellular proteome. Science (New York, N.Y.), 320(5877), 807–811. 10.1126/science.1154370

35. Liu, C., Li, X., Huang, Q., Zhang, M., Lei, T., Wang, F., Zou, W., Huang, R., Hu, X., Wang, C., Zhang, X., Sun, B., Xing, L., Yue, J., & Yu, J. (2023). Single-cell RNA-sequencing reveals radiochemotherapy-induced innate immune activation and MHC-II upregulation in cervical cancer. Signal transduction and targeted therapy, 8(1), 44. 10.1038/s41392-022-01264-9

36. Liu, C., Zhang, M., Yan, X., Ni, Y., Gong, Y., Wang, C., Zhang, X., Wan, L., Yang, H., Ge, C., Li, Y., Zou, W., Huang, R., Li, X., Sun, B., Liu, B., Yue, J., & Yu, J. (2023). Single-cell dissection of cellular and molecular features underlying human cervical squamous cell carcinoma initiation and progression. Science advances, 9(4), eadd8977. 10.1126/sciadv.add8977

37. Lundberg, M., Leivo, I., Saarilahti, K., Mäkitie, A. A., & Mattila, P. S. (2012). Transforming growth factor beta 1 genotype and p16 as prognostic factors in head and neck squamous cell carcinoma. Acta Oto-Laryngologica, 132(9), 1006–1012. 10.3109/00016489.2012.678944

38. Maarifi, G., Czubala, M. A., Lagisquet, J., Ivory, M. O., Fuchs, K., Papin, L., Birchall, J. C., Nisole, S., Piguet, V., & Blanchet, F. P. (2020). Langerin (CD207) represents a novel interferon-stimulated gene in Langerhans cells. Cellular & molecular immunology, 17(5), 547–549. 10.1038/s41423-019-0302-5

39. Malagón, T., Franco, E. L., Tejada, R., et al. (2024). Epidemiology of HPV-associated cancers past, present and future: Towards prevention and elimination. *Nature Reviews Clinical Oncology, 21*, 522–538. 10.1038/s41571-024-00904-z

40. McCarthy, D. J., Campbell, K. R., Lun, A. T. L., & Willis, Q. F. (2017). Scater: Pre-processing, quality control, normalisation and visualisation of single-cell RNA-seq data in R. *Bioinformatics, 33*, 1179-1186. 10.1093/bioinformatics/btw777

41. Mizumoto, N., & Takashima, A. (2004). CD1a and langerin: acting as more than Langerhans cell markers. The Journal of clinical investigation, 113(5), 658–660. 10.1172/JCI21140

42. Moon, H. S., Park, W. I., Sung, S. H., Choi, E. A., Chung, H. W., & Woo, B. H. (2003). Immunohistochemical and quantitative competitive PCR analyses of midkine and pleiotrophin expression in cervical cancer. Gynecologic oncology, 88(3), 289–297. 10.1016/s0090-8258(02)00070-7

43. Moraes, L. A., Kar, S., Foo, S. L., et al. (2017). Annexin-A1 enhances breast cancer growth and migration by promoting alternative macrophage polarization in the tumour microenvironment. Scientific Reports, 7(1), 17925. 10.1038/s41598-017-17622-5

44. Ogawa, S., Matsuoka, Y., Takada, M., Matsui, K., Yamane, F., Kubota, E., Yasuhara, S., Hieda, K., Kanayama, N., Hatano, N., Tokumitsu, H., & Magari, M. (2019). Interleukin 34 (IL-34) cell-surface localization regulated by the molecular chaperone 78-kDa glucose-regulated protein facilitates the differentiation of monocytic cells. The Journal of biological chemistry, 294(7), 2386–2396. 10.1074/jbc.RA118.006226

45. Oto, J., Le, Q.-K., Schäfer, S. D., Kiesel, L., Marí-Alexandre, J., Gilabert-Estellés, J., Medina, P., & Götte, M. (2023). Role of syndecans in ovarian cancer: New diagnostic and prognostic biomarkers and potential therapeutic targets. *Cancers, 15*(12), 3125. 10.3390/cancers15123125

46. Otsuka, R., Wada, H., & Seino, K. I. (2021). IL-34, the rationale for its expression in physiological and pathological conditions. Seminars in immunology, 54, 101517. 10.1016/j.smim.2021.101517

47. Ou, Z., Lin, S., Qiu, J., Ding, W., Ren, P., Chen, D., Wang, J., Tong, Y., Wu, D., Chen, A., Deng, Y., Cheng, M., Peng, T., Lu, H., Yang, H., Wang, J., Jin, X., Ding, M., Xu, X., … Wu, P. (2022). Single-nucleus RNA sequencing and spatial transcriptomics reveal the immunological microenvironment of cervical squamous cell carcinoma. *bioRxiv*. 10.1101/2021.12.23.473944

48. Özbay Kurt, F. G., Cicortas, B. A., Balzasch, B. M., De la Torre, C., Ast, V., Tavukcuoglu, E., Ak, C., Wohlfeil, S. A., Cerwenka, A., Utikal, J., & Umansky, V. (2024). S100A9 and HMGB1 orchestrate MDSC-mediated immunosuppression in melanoma through TLR4 signaling. Journal for immunotherapy of cancer, 12(9), e009552. 10.1136/jitc-2024-009552

49. Pham, D., Tan, X., Balderson, B. et al. Robust mapping of spatiotemporal trajectories and cell– cell interactions in healthy and diseased tissues. Nat Commun 14, 7739 (2023). 10.1038/s41467-023-43120-6

50. Peyre, L., Meyer, M., Hofman, P., & Roux, J. (2021). TRAIL receptor-induced features of epithelial-to-mesenchymal transition increase tumour phenotypic heterogeneity: potential cell survival mechanisms. British journal of cancer, 124(1), 91–101. 10.1038/s41416-020-01177-w

51. Pinto, A., Schlecht, N., Woo, T. Y. C., et al. (2008). Biomarker (ProExTM C, p16INK4A, and MiB-1) distinction of high-grade squamous intraepithelial lesion from its mimics. *Modern Pathology, 21*, 1067–1074. 10.1038/modpathol.2008.101

52. Rettig, E. M., Zaidi, M., Faraji, F., Eisele, D. W., El Asmar, M., Fung, N., D’Souza, G., & Fakhry, C. (2018). Oropharyngeal cancer is no longer a disease of younger patients and the prognostic advantage of Human Papillomavirus is attenuated among older patients: Analysis of the National Cancer Database. Oral oncology, 83, 147–153. 10.1016/j.oraloncology.2018.06.013

53. Ritchie, M. E., Phipson, B., Wu, D., Hu, Y., Law, C. W., Shi, W., & Smyth, G. K. (2015). limma powers differential expression analyses for RNA-sequencing and microarray studies. *Nucleic Acids Research, 43*(7), e47. 10.1093/nar/gkv007

54. Singh, D., Vignat, J., Lorenzoni, V., Eslahi, M., Ginsburg, O., Lauby-Secretan, B., Arbyn, M., Basu, P., Bray, F., & Vaccarella, S. (2023). Global estimates of incidence and mortality of cervical cancer in 2020: A baseline analysis of the WHO Global Cervical Cancer Elimination Initiative. *The Lancet Global Health, 11*(2), e197–e206. 10.1016/S2214-109X(22)00501-0

55. Singh, M., Mehrotra, S., Kalra, N., Singh, U., & Shukla, Y. (2008). Correlation of DNA ploidy with progression of cervical cancer. *Journal of Cancer Epidemiology, 2008*, 298495. 10.1155/2008/298495

56. Singini, M. G., Muchengeti, M., Sitas, F., Chen, W. C., Combes, J. D., Waterboer, T., & Clifford, G. M. (2024). Antibodies against high-risk human papillomavirus proteins as markers for noncervical HPV-related cancers in a Black South African population, according to HIV status. International journal of cancer, 155(2), 251–260.

57. Strachan, D. C., Ruffell, B., Oei, Y., Bissell, M. J., Coussens, L. M., Pryer, N., & Daniel, D. (2013). CSF1R inhibition delays cervical and mammary tumor growth in murine models by attenuating the turnover of tumor-associated macrophages and enhancing infiltration by CD8+ T cells. Oncoimmunology, 2(12), e26968. 10.4161/onci.26968

58. Stuart, T., Butler, A., Hoffman, P., Hafemeister, C., Papalexi, E., Mauck, W. M., 3rd, Hao, Y., Stoeckius, M., Smibert, P., & Satija, R. (2019). Comprehensive integration of single-cell data. *Cell, 177*(7), 1888–1902.e21. 10.1016/j.cell.2019.05.031

59. Sun, L., Su, Y., Jiao, A., Wang, X., & Zhang, B. (2023). T cells in health and disease. Signal transduction and targeted therapy, 8(1), 235. 10.1038/s41392-023-01471-y

60. Supek, F., Bošnjak, M., Škunca, N., & Šmuc, T. (2011). REVIGO summarizes and visualizes long lists of gene ontology terms. *PloS One, 6*(7), e21800. 10.1371/journal.pone.0021800

61. Syrjänen, S., Naud, P., Sarian, L., Derchain, S., Roteli-Martins, C., Longatto-Filho, A., Tatti, S., Branca, M., Erzen, M., Hammes, L. S., Costa, S., & Syrjänen, K. (2009). Up-regulation of plasminogen activator inhibitor-2 is associated with high-risk HPV and grade of cervical lesion at baseline but does not predict outcomes of high-risk HPV infections or incident CIN. American journal of clinical pathology, 132(6), 883–892. 10.1309/AJCPQQ07WUTZUTES

62. Tuong, Z. K., Lukowski, S. W., Nguyen, Q. H., Chandra, J., Zhou, C., Gillinder, K., Bashaw, A. A., Ferdinand, J. R., Stewart, B. J., Teoh, S. M., Hanson, S. J., Devitt, K., Clatworthy, M. R., Powell, J. E., & Frazer, I. H. (2021). A model of impaired Langerhans cell maturation associated with HPV induced epithelial hyperplasia. iScience, 24(11), 103326. 10.1016/j.isci.2021.103326

63. Varga, N., Mózes, J., Keegan, H., White, C., Kelly, L., Pilkington, L., Benczik, M., Zsuzsanna, S., Sobel, G., Koiss, R., Babarczi, E., Nyíri, M., Kovács, L., Attila, S., Kaltenecker, B., Géresi, A., Kocsis, A., O’Leary, J., Martin, C. M., & Jeney, C. (2017). The Value of a Novel Panel of Cervical Cancer Biomarkers for Triage of HPV Positive Patients and for Detecting Disease Progression. Pathology oncology research : POR, 23(2), 295–305. 10.1007/s12253-016-0094-1

64. Wang, X., Huang, X., & Zhang, Y. (2018). Involvement of human papillomaviruses in cervical cancer. *Frontiers in Microbiology, 9*, 2896. 10.3389/fmicb.2018.02896

65. Wang, Y., Szretter, K., Vermi, W. et al. IL-34 is a tissue-restricted ligand of CSF1R required for the development of Langerhans cells and microglia. Nat Immunol 13, 753–760 (2012). 10.1038/ni.2360

66. Xie, Z., Bailey, A., Kuleshov, M. V., Clarke, D. J. B., Evangelista, J. E., Jenkins, S. L., Lachmann, A., Wojciechowicz, M. L., Kropiwnicki, E., Jagodnik, K. M., Jeon, M., & Ma’ayan, A. (2021). Gene set knowledge discovery with Enrichr. *Current Protocols, 1*, e90. 10.1002/cpz1.90

67. Yano, H., Andrews, L. P., Workman, C. J., & Vignali, D. A. A. (2019). Intratumoral regulatory T cells: markers, subsets and their impact on anti-tumor immunity. Immunology, 157(3), 232–247. 10.1111/imm.13067

68. Zhang, J., Zhang, Y., & Zhang, Z. (2018). Prevalence of human papillomavirus and its prognostic value in vulvar cancer: A systematic review and meta-analysis. PloS one, 13(9), e0204162. 10.1371/journal.pone.0204162

