## Supplementary Tables for "Spatial analysis of HPV associated cervical intraepithelial neoplastic tissues demonstrate distinct immune signatures associated with cervical cancer progression"

| Immune Cell Type | Marker Genes |
| --- | --- |
| T-regulatory | "CD4", "TNFRSF18", "IL2RA", "FOXP3", "TGFB1", "IL10" |
| T pro-inflammatory | "CSF3","IL1A","IL1B","IL6","IFNG", "IL4", "IL5", "IL13", "IL36B","IL36G","IL36A", "CXCL8", "TNF", "IL18" |
| T anti-inflammatory | "IL10","IL12A","IL12B", "IL22", "IL37", "IL1F10", "TGFB1","IL4", "IL11", "IL13" |
| M1 macrophages | ("CD80", "IFNG","IL1B", "IL6", "TNF", "CCL2", "FCGR3A", "FCGR2A", "FCGR1A", "IL12B", "IL23A", "MARCO", "CD86" |
| M2 macrophages | "VTCN1","CD36","CD200R1","CD163","MRC1","CD209","CLEC10A ","CLEC7A","CXCR1", "CXCR2","MSR1", "ARG1","NOS2", "RETNLB","HMOX1", "PPARG","CD274", "IL1RN", "TGM2", "ACTG1", "TIMP1", "SPHK1", "CXCL1", "MERTK", "ITGAV", "IL1A", "VEGFA", "IVNS1ABP", "CXCL8", "DYSF", "EHHADH", "RIF1", "LIG4" |
| Langerhans | "CD207", "CD1A" |

**Supplementary Table 1**. List of all marker genes used for immune cell type classification

| Module Score Groups | Marker Genes |
| --- | --- |
| IL34-CSF1R | “IL34”, “CSF1R” |
| Cervical Cancer | "SERPINB3", "TP63", "KRT5", "CDKN2A" |

**Supplementary Table 2**. List of all marker genes used to identify average co-expression values.
