## Supplementary Figures for "Spatial analysis of HPV associated cervical intraepithelial neoplastic tissues demonstrate distinct immune signatures associated with cervical cancer progression"

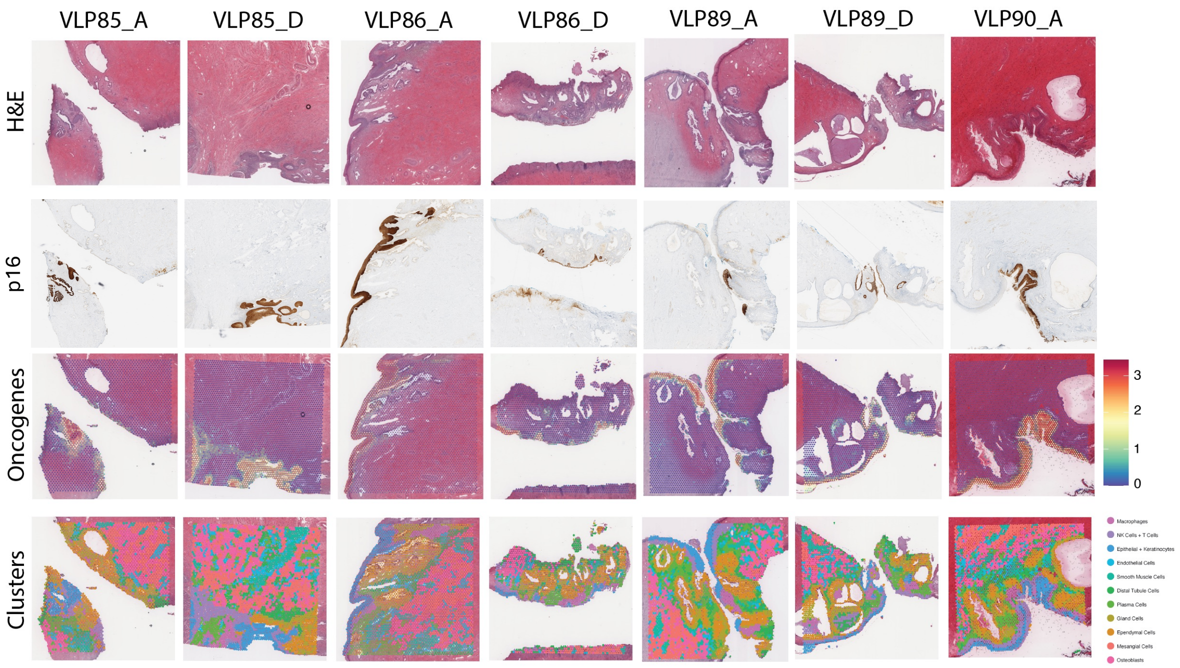


**Supplementary Fig 1.** H&E image, p16 immunohistochemistry staining, spatial clustering, and co-expression of cervical cancer oncogenes (*CDKN2A*, *SERPINB3*, *TP63* and *KRT5*) for all seven samples.

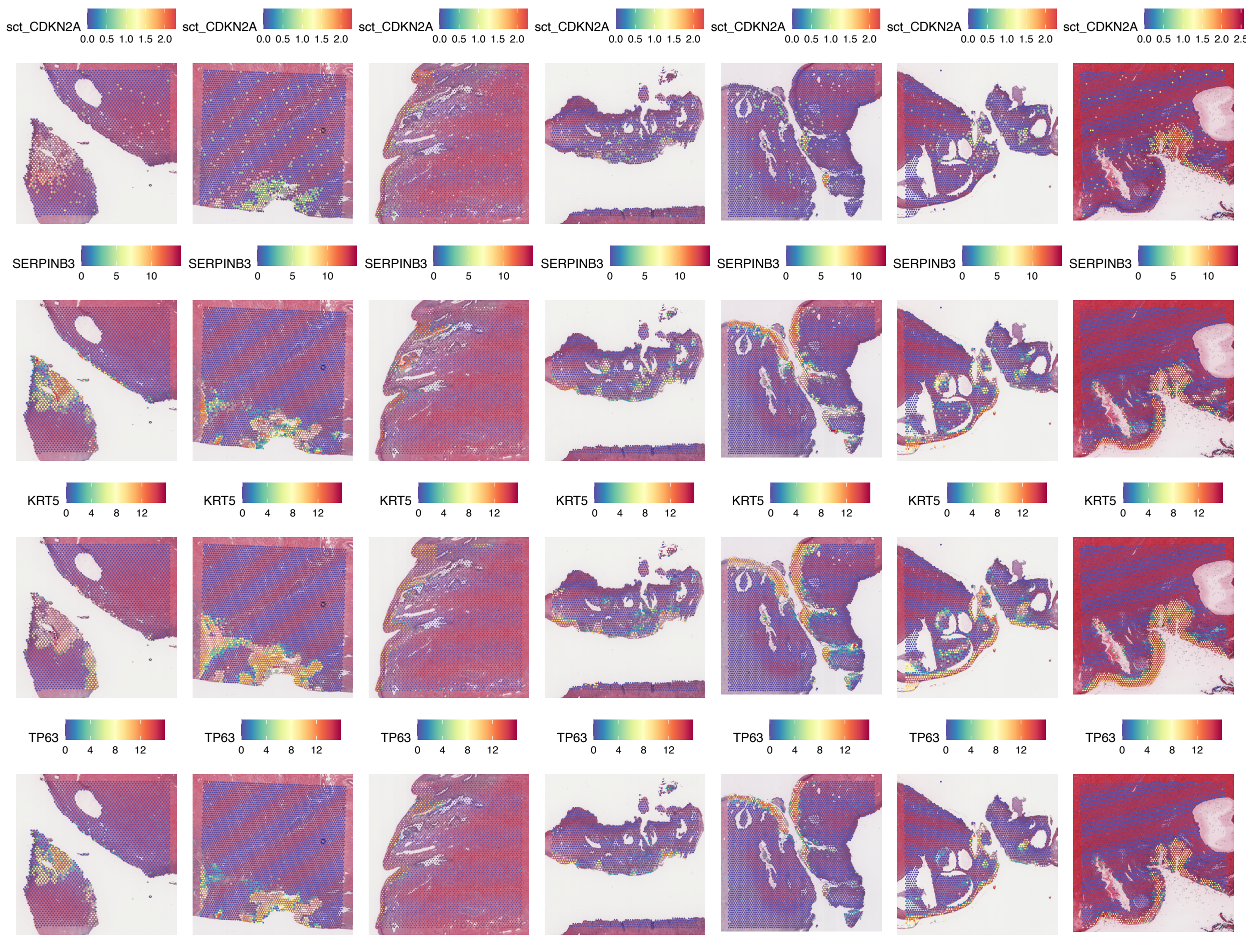


**Supplementary Fig 2.** Spatial gene expression of cervical cancer oncogenes (*CDKN2A*, *SERPINB3*, *KRT5* and *TP63*) for all seven samples.

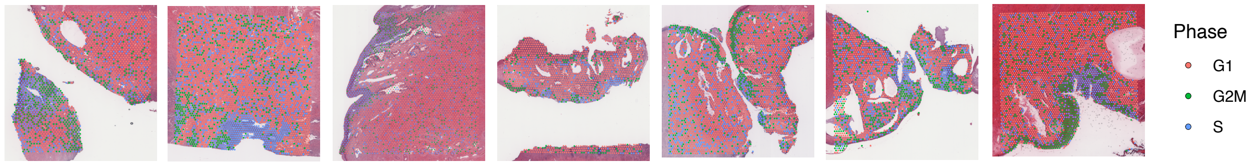


**Supplementary Fig 3.** Cell cycle scoring analysis across all seven samples.

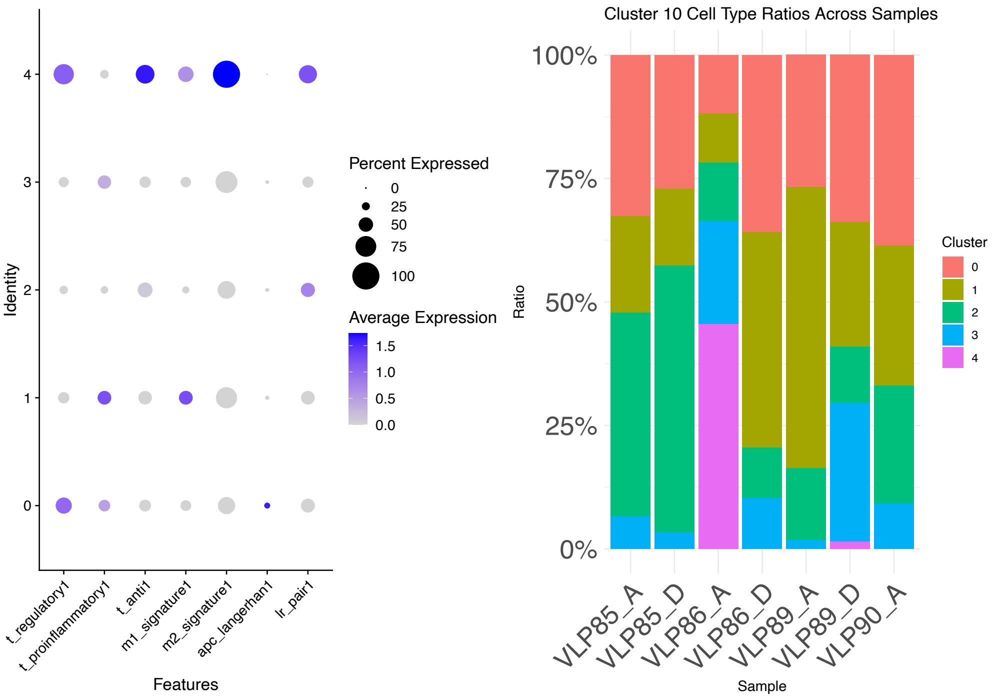


**Supplementary Fig 4.** Sub-clustering of cluster 10: Immune cell type expression and cell type proportions for each sample. Clusters 0 - 4 representative of sub-clusters within cluster 10.

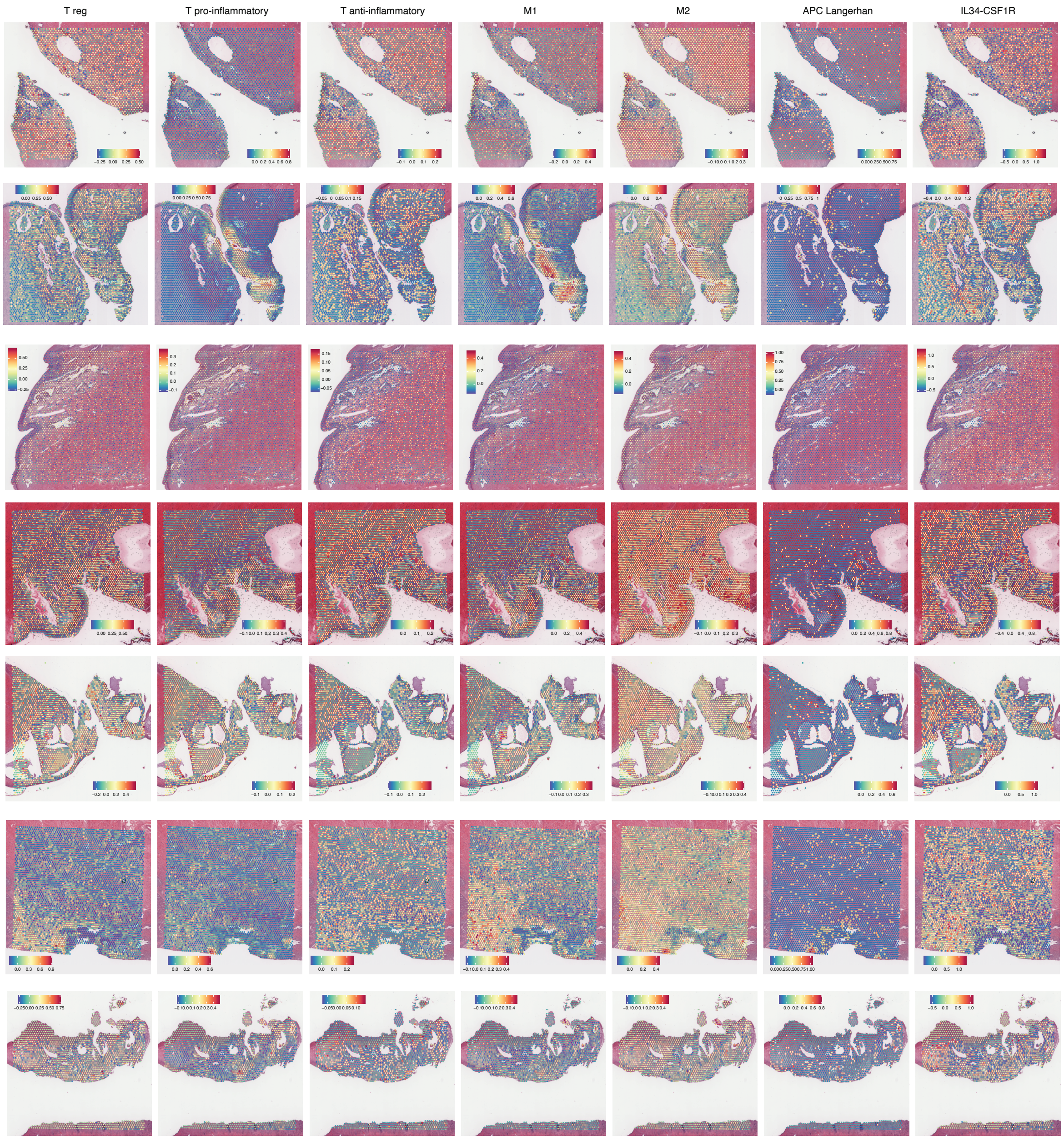


**Supplementary Fig 5.** Spatial mapping of immune cell types; T regulatory, T pro-inflammatory, T anti-inflammatory, M1 macrophages, M2 macrophages, Langerhans cells, and IL34-CSF1R co-expression, from left to right for all seven samples.

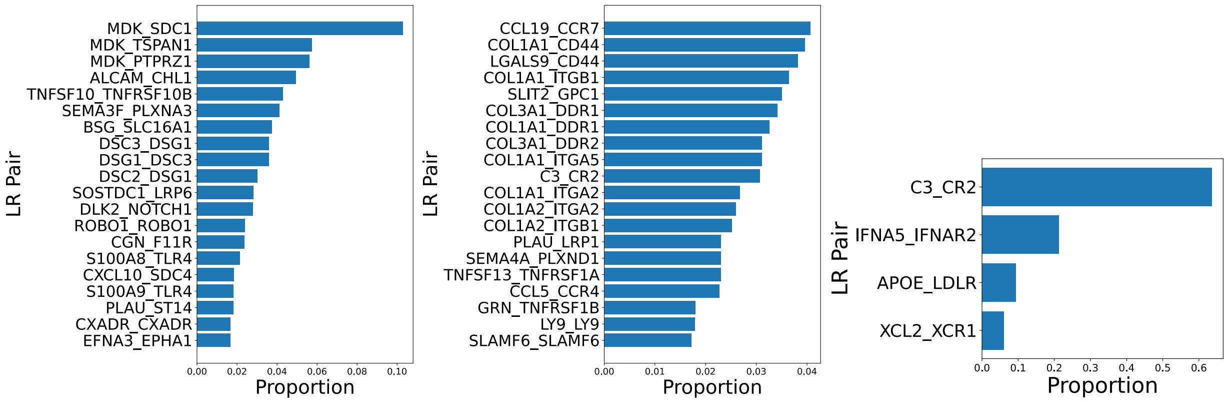


**Supplementary Fig 6.** Top 20 interacting ligand-receptor pairs shown between clusters 8 and 9 (left) and clusters 9 and 10 (middle). All ligand-receptor interactions are shown between clusters 8 and 10 (right).
